# Embracing curiosity eliminates the exploration-exploitation dilemma

**DOI:** 10.1101/671362

**Authors:** Erik J Peterson, Timothy D Verstynen

## Abstract

Balancing exploration with exploitation is seen as a mathematically intractable dilemma that all animals face. In this paper, we provide an alternative view of this classic problem that does not depend on exploring to optimize for reward. We argue that the goal of exploration should be pure curiosity, or learning for learning’s sake. Through theory and simulations we prove that explore-exploit problems based on this can be solved by a simple rule that yields optimal solutions: when information is more valuable than rewards, be curious, otherwise seek rewards. We show that this rule performs well and robustly under naturalistic constraints. We suggest three criteria can be used to distinguish our approach from other theories.

## Introduction

The exploration-exploitation dilemma is seen as a fundamental, yet intractable, problem in the biological world (1–6). In this problem, the actions taken are based on a set of learned values (3, 7). Exploitation is defined as choosing the most valuable action. Exploration is defined as making any other choice. Exploration and exploitation are thus seen as opposites, but their trade-off is not formally a dilemma.

To create the dilemma two more things must be true. First, is that exploration and exploitation have a shared objective. This is satisfied in reinforcement learning when exploration and exploitation both try to maximize rewards collected from the environment (3). Second uncertainty about the outcome must be present prior to action (3). Combined these two assumptions create and define the classic exploration-exploitation dilemma, which we illustrate in Figure 1a. They also make a tractable solution to the problem impossible (2, 4, 5).

**Fig. 1.**
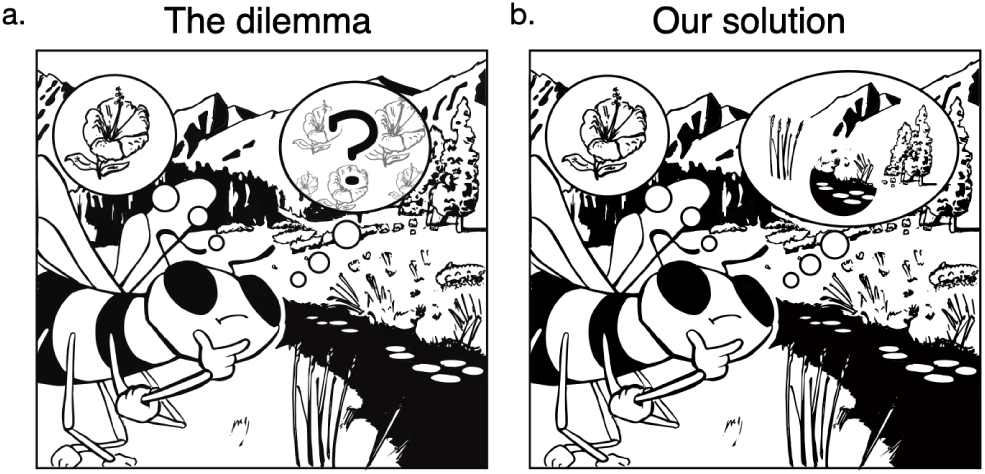
Two views of exploration and exploitation from the point of view of a bee. **a**. The classic dilemma: either exploit an action with a known reward by returning to the best flower or explore other flowers on the chance they will return a better outcome. The central challenge here is that the exploration of other flowers is innately uncertain in terms of the pollen collected, the extrinsic reward. **b**. Our alternative: a bee can have two goals: either exploit the pollen from the best flower, *or* explore to maximize pure learning with intrinsically curious search of the environment. The solution we propose to our alternative is the bee or any other learning agent ought to pursue either intrinsic information value or extrinsic reward value, depending on which is expected to be larger. In other words, to solve explore-exploit decisions we set information and reward on equal terms, and propose a greedy rule to choose between them. *Artist credit*: Richard Grant.

The dilemma in this form has been studied for decades and a variety of complex and approximate answers have been prooposed (2, 5, 6, 8–14). We put aside this work in order to ask a critical and contrarian question: *is the dilemma really a fundamental problem for animals to solve?*

In this paper we offer an alternative view. We hypothesize that curiosity is a sufficient motivation for all exploration. We then treat extrinsic rewards and intrinsic curiosity as equally important, but independent motivations (15). We put them in direct competition with each other, as illustrated in Figure 1b.

Ther is no one reason to suppose curiosity is sufficient, there are many. Curiosity leads to the building of intuitive physics (16) and is key to understanding causality (17). It can help ensure that an animal has generalizable world models (18, 19). It can drive imagination, play, and creativity (20–23). It can lead to the discovery of, and creation of, knowledge (24–26). It can ensure local minima, or other deceptive outcomes, are overcome when learning (27, 28). It leads to the discovery of changes in the environment (29), and ensures there are robust action policies to respond to these changes (30). It can help in the recovery from injury (31). It can drive language learning (32). It is critical to evolution and the process of cognitive development (33, 34). Animals will prefer curiosity over extrinsic rewards, even when that information is costly (35–38). It is this combination of reasons that makes curiosity useful to an organism and it is this broad usefulness that let’s us hypothesize that curiosity is sufficient for all exploration (15, 27, 39).

Our simple approach is a stronger view than is standard. Normally, to solve dilemma problems, exploration is motivated by extrinsic rewards and aided by “breadcrumbs” from intrinsic rewards (9, 40, 41). Approximating a dilemma solution is normally about choosing the algorithm that creates these “breadcrumbs”, and then setting their relative importance compared to environmental rewards. Instead, we contrast intrinsically motivated exploration with pure extrinsic exploitation in a competitive game that is played out inside the minds of individual animals.

This paper consists of a new union of pure intrinsic and pure extrinsic motivations. We prove there is an optimal value rule to choose between curiosity and environmental rewards. We follow this up with naturalistic simulations, showing where our union’s performance matches or exceeds normal approaches, and we show that overall pure exploration is the more robust solution to the dilemma of choosing to explore or chooosing to exploit.

## Results

### Reward collection – theory

When the environment is unknown, an animal needs to trade-off exploration and exploitation. The standard way of approaching this is to assume, “The agent must try a variety of actions and progressively favor those that appear to be the most rewarding” (3).

We first consider an animal who interacts with an environment over a sequence of states, actions and rewards. The animal’s goal is to select actions in a way that maximizes the total reward collected. How should an animal do this?

To describe the environment we use a discrete time Markov decision process, 𝒳_*t*_ = (𝒮, 𝒜, 𝒯, ℛ). Specifically, states are real valued vectors *S* from a finite set of 𝒮 size *n*. Actions *A* are from the finite real set 𝒜 of size *k*. We consider a set of policies that consist of a sequence of functions, *π* = {*π*_*t*_, *π*_*t*+1_, …, *π*_*T*−1_}. For any step *t*, actions are generated from states by policies, either deterministic *A* = *π*(*S*) or random *A* ∼ *π*(*S* |*A*). In our presentation we drop the indexing notation on our policies, using simply *π* to refer to the sequence as a whole. Rewards are single valued real numbers, *R*, generated by a reward function *R* ∼ ℛ(*S*|*A, t*). Transitions from state *S* to new state *S*t′ are caused by a stochastic transition function, *S*′ ∼ 𝒯(*S*|*A, t*). We leave open the possibility both 𝒯 and ℛ may be time-varying.

In general we use the = to denote assignment, in contrast to ∼ which we use to denote taking random samples from some distribution. The standalone | operator is used to denote conditional probability. An asterisk is used to denote optimal value policies, *π*\*.

We focus on finite time horizons. We do not discount the rewards. Both for simplicity. Our basic approach should generalize to continuous spaces, and discounted but infinite time horizons (42)

So given a policy *π*_*R*_, the value function in standard reinforcement learning is given by Eq. 1. This the term we use to maximize reward collection.

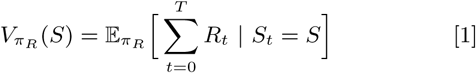

But this equation is hard to use in practice because it requires we integrate over all time, *T*. The Bellman equation (Eq. 2) is a desirable simplification because it reduces the entire action sequence in Eq. 1 into two (recursive) steps, *t* and *t* + 1. These steps can then be recursively applied.

The practical problem we are with left in a this simplification is finding a reliable estimate of 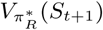 (Eq. 2). This is a problem we return to further on.

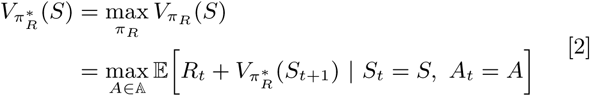

### Information collection – theory

Should any one role for curiosity dominate the others? We suggest no. We therefore define a new metric of *learning progress* (43) that is general, practical, simple, and compatible with the normal limits of experimental knowledge. That is, we do not assume the learning algorithm or goal is known. We only assume there is a memory whose dynamics we can measure.

We now consider an animal who interacts with an environment over the same space as in reward collection, but whose goal is to select actions in a way that maximizes information value (26, 44–52). In this, we face two questions: How should this information be valued and collected over time? And like for reward collection, is there a Bellman solution?

We assume a Bellman solution is possible and set about finding it. Let’s then say *E* represents the information value. The Bellman solution we want for *E* is given by Eq. 3.

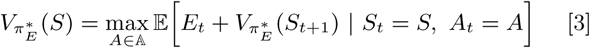

We assume that maximizing *E* also maximizes curiosity. We refer to our deterministic approach to curiosity as E-exploration. Or E-explore, for short. In the following paragraphs we will define *E*, and prove this definition has the Bellman solution shown in Eq.3. We will *not* limit ourselves to Markovian ideas of learning and memory in doing this.

We define memory **M** as a vector of size *p* embedded in a real-valued space that is closed and bounded. This idea maps to a range of physical examples of memory, including firing rates of neurons in a population (53), strength differences between synapses (54), or calcium dynamics (55).

A learning function is then any function *f* that maps observations 𝒳 into memory **M**. This idea can be expressed using recursive notation, denoted by **M** ← *f* (𝒳, **M**). Though we will sometimes use **M**′ to denote the updated memory instead. We also need to define a forgetting function, *f* ^−1^*f* (𝒳, **M**′) → **M** which is important later. As an auxiliary assumption we assume *f* has been selected by evolution to be formally learnable (56).

### Curiosity axioms

Should any one role for curiosity dominate the others? We suggest no. We therefore define a new metric of *learning progress* (43) that is general, practical, simple, and compatible with the normal limits of experimental measurements. That is, we do not assume the learning algorithm, or target goal, are known. We only assume we have a memory, who’s learning dynamics we can measure.

We reason the value of any information depends entirely on how much memory changes in making an observation. We formalize this idea with axioms, which follow.

These axioms are useful in three ways: First, they give a measure of value information *without* needing to know the exact learning algorithm(s) in use. This is helpful because we rarely know how an animal is learning in detail. This leads to the second useful property. If we want to measure a subjects intrinsic information value, we need only record differences in dynamics of their memory circuits. There is little need to decode, or interpret. Third, these definitions properly generalize all prior attempts to formalize curiosity.

So if we have a memory **M** which has been learned by *f* over a path of observations, (**X**_**0**_, **X**_**1**_, …), can we measure how much value *E* the next observation **X**_**t**_ should have?

#### Axiom 1 (Axiom of Memory).

*E is continuous with continuous changes in memory*, Δ**M**, *between* **M**′ *and f* (𝒳, **M**).

#### Axiom 2 (Axiom of Specificity).

*If all p elements* |Δ*M*_*i*_| *in* **M** *are equal, then E is minimized*.

That is, information that is learned evenly in memory cannot induce any inductive bias, and is therefore nonspecific and is the least valuable learning possible (57).

#### Axiom 3 (Axiom of Scholarship).

*Learning has an innately positive value, even if later on it has some negative consequences. Therefore, E* ≥ 0.

#### Axiom 4 (Axiom of Equilibrium).

*Given the same observation* 𝒳_*i*_, *learning will reach equilibrium. Therefore E will decrease to below some finite threshold η >* 0 *in some finite time T*.

See Figure 2a for an example of *E* defined in memory space. We show examples of learning equilibrium and (self-)consistent and inconsistent learning in Figure 2b-c.

**Fig. 2.**
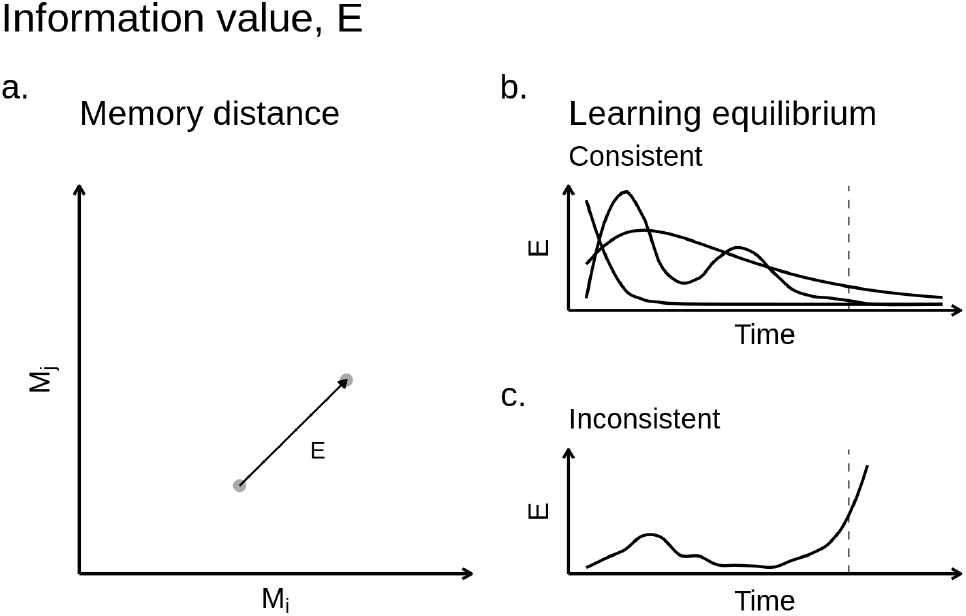
Information value. **a**. This panel illustrates a two dimensional memory. The information value of an observation depends on the distance memory “moves” during learning, denoted here by *E* We depict memory distance as euclidean space, but this is one of many possible ways to realize *E*. **b-c** This panel illustrates some examples of learning dynamics in time, made over a series of observations that are not shown. If information value becomes decelerating in finite time bound, then we say learning is consistent with Def. 2. This is what is shown in panel b. If learning does not decelerate, then it is said to be inconsistent (Panel c). The finite bound is represented here by a dotted line.

Note that a corollary of Axiom 1 is that an observation that doesn’t change memory, has no value. So if 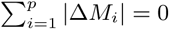 then *E* = 0. It also follows from Axiom 3 and 4 that E-exploration will visit every state in 𝒮 ar least once in a finite time *T*, assuming *E*_0_ *>* 0. And as we stop exploration when *E < η*, only states in which there is more to be learned will be revisited. This, along with our deterministic policy, ensures perfectly efficient exploration in terms of learning progress.

### A Bellman solution

A common way to arrive at the Bellman solution is to rely on a Markov decision space. This is a problem for our definition of memory as it has path dependence and so violates the Markov property. To get around this, we prove that exact forgetting of the last observation is another way to find a Bellman solution. Having proven the optimal substructure of **M** (see the Appendix), and assuming some arbitrary starting value *E*_0_ *>* 0, it follows that the Bellman solution to information optimization is in fact given by Equation 3.

### The importance of boredom

To limit curiosity–to avoid the white noise problem (18), other useless minutia (58), and to stop exploration efficiently–we rely on the threshold term, *η* ≥ 0 (Ax. 4). We treat *η* as synonymous with boredom (9, 59, 60). In other words, we hypothesize boredom is an adaptive parameter tuned to fit the environment (61). Specifically, we use boredom to ignore arbitrarily small amounts of information value, by requiring exploration of some observation 𝒳 to cease once *E* ≤ *η* for that 𝒳.

### Reward and information – theory

Finally, our union of independent curiosity with independent reward collection. We now model an animal who interacts with an environment who wishes to maximize both information *and* reward value, as shown in Eq 4. How can an animal do this? To answer this, let’s make two conjectures:

**Conjecture 1** (The hypothesis of equal importance). *Reward and information collection are equally important for survival*.

**Conjecture 2** (The curiosity trick). *Curiosity is a sufficient solution for all exploration problems (where learning is possible)*.

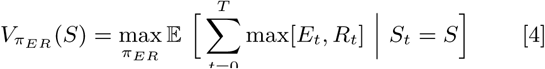

Having already established we have a Bellman optimal policy for *E*, and knowing reinforcement learning provides many solutions for *R* (3, 42), we can write out the Bellman optimal solution for 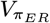. This is,

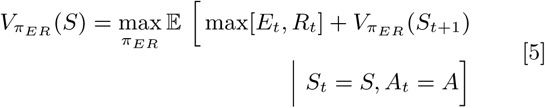

From here we can substitute in the respective value functions for reward and information value. This gives us Eq.6, which translates to the Bellman optimal decision policy shown in Eq.7.

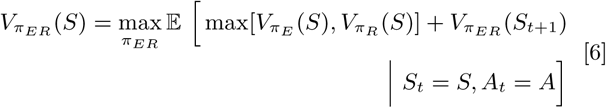

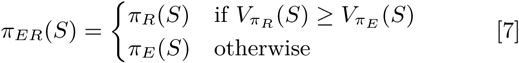

### Simplifying with win-stay lose-switch

In our analysis of decision making we have assumed value for the next state, *V* (*S*_*t*+1_) is readily and accurately available. In practice for most animals this is often not so, and is a key distinction between the field of dynamic programming and the more biologically sound reinforcement learning. If we no longer assume 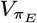 and 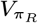 are available but must be estimated from the environment, then we are left with something of a paradox.

We wish to use value functions to decide between information and reward value but are simultaneously estimating those values. There are a range of methods to handle this (3, 42), but we opted to further simplify *π*_*ER*_ in three ways. We first shifted from using the full value functions to using only the last payout, *E*_*t*−1_ and *R*_*t*−1_. Second, we removed all state-dependence leaving only time dependence. Third, we also included *η*0 to control the duration of exploration. This gives our final working equation (Eq. 8 a new sort of win-stay lose-switch (WSLS) rule.

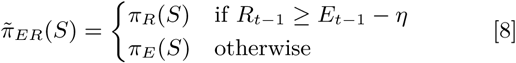

In Eq. 8, we think of *π*_*R*_ and *π*_*E*_ as two “players” in a game played for behavioral control (62). We feel the approach has several advantages. It is myopic and therefore can optimally handle nonlinear changes in either the reward function or environmental dynamics (63). In stationary settings, its regret is bounded (64). It can approximate Bayesian inference (65). It leads to cooperative behavior (66). Further, WSLS has a long history in psychology, where it predicts and describes human and animal behavior (67). But most of all, our version of WSLS is simple to implement, but robust in practice (shown below). It should therefore scale well in different environments and species.

### Information collection – simulations

The optimal policy to collect information value is not a sampling policy. It is a deterministic greedy maximization (Eq 3). In other words, to minimize uncertainty during exploration an animal should not introduce additional uncertainty by taking random actions (68).

To confirm our deterministic method is best we examined a multi-armed information task (Task 1; Figure 3a). This variation of a bandit task replaced rewards with information. In this case, simple colors. On each selection, the agent saw one of two colors (integers) returned according to specific probabilities (Figure 3a).

**Fig. 3.**
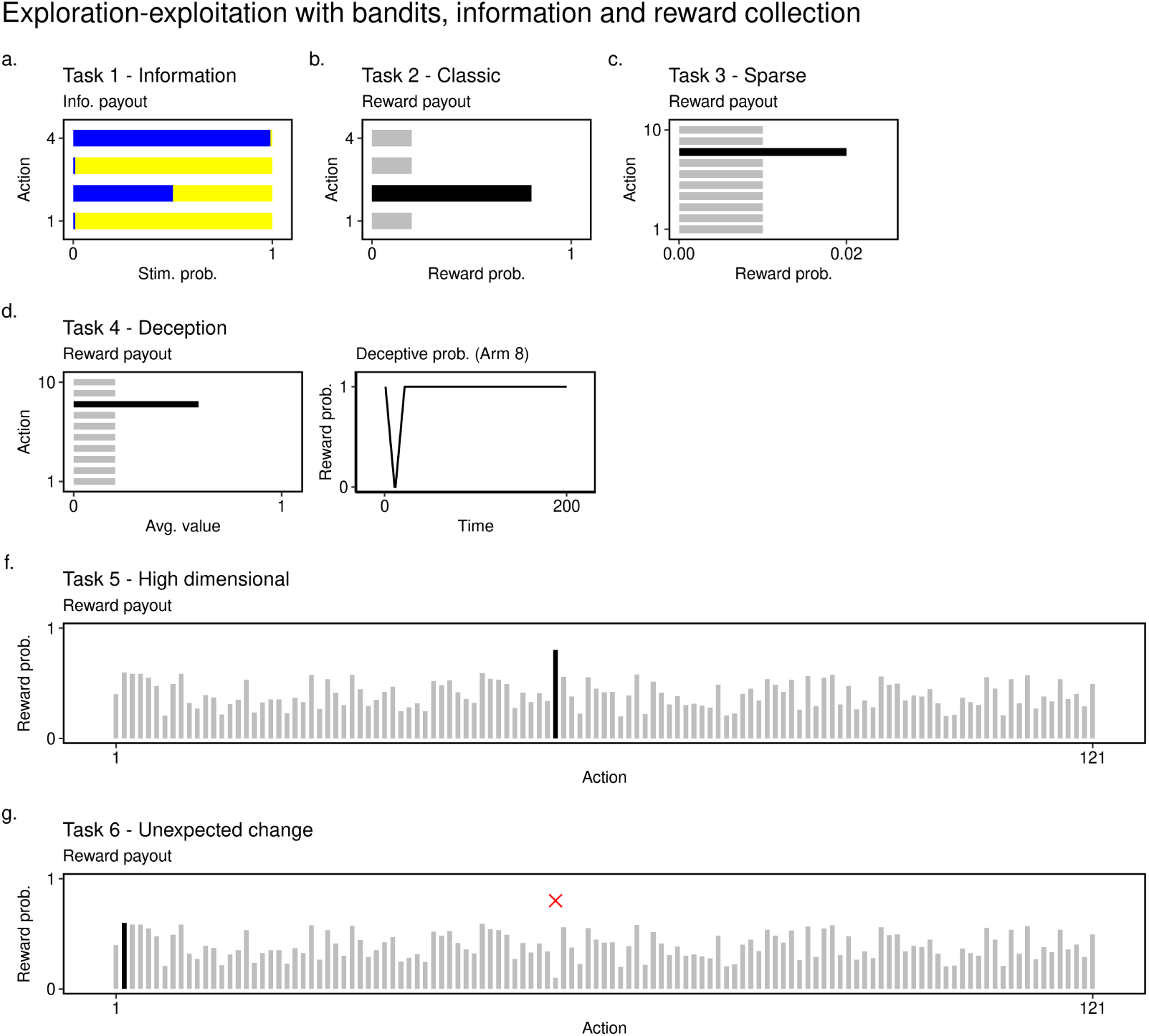
Multi-armed bandits – illustration and payouts. On each trial the agent must take on *n* actions. Each action generates a payout. Payouts can be information, reward, or both. For comments on general task design. **a**. A 4 choice bandit for information collection. In this task the payout is information, a yellow or blue “stimulus”. A good agent should visit each arm, but quickly discover that only arm two is information bearing. **b**. A 4 choice design for reward collection. The agent is presented with four actions and it must discover which choice yields the highest average reward. In this task that is Choice 2. **c**. A 10 choice sparse reward task. Note the very low overall rate of rewards. Solving this task with consistency means consistent exploration. **d**. A 10 choice deceptive reward task. The agent is presented with 10 choices but the action which is the best in the long-term (>30 trials) has a lower value in the short term. This value first declines, then rises (see column 2). **e**. A 121 choice task with a complex payout structure. This task is thought to be at the limit of human performance. A good agent will eventually discover choice number 57 has the highest payout. **f**. This task is identical to *a*., except for the high payout choice being changed to be the lowest possible payout. This task tests how well different exploration strategies adjust to simple but sudden change in the environment.

In Figure 4 we compared optimal value E-exploration to a noisy version of itself. As predicted, determinism generated more value in less time, when compared against a (slightly) stochastic agent using an otherwise identical curiosity-directed search.

**Fig. 4.**
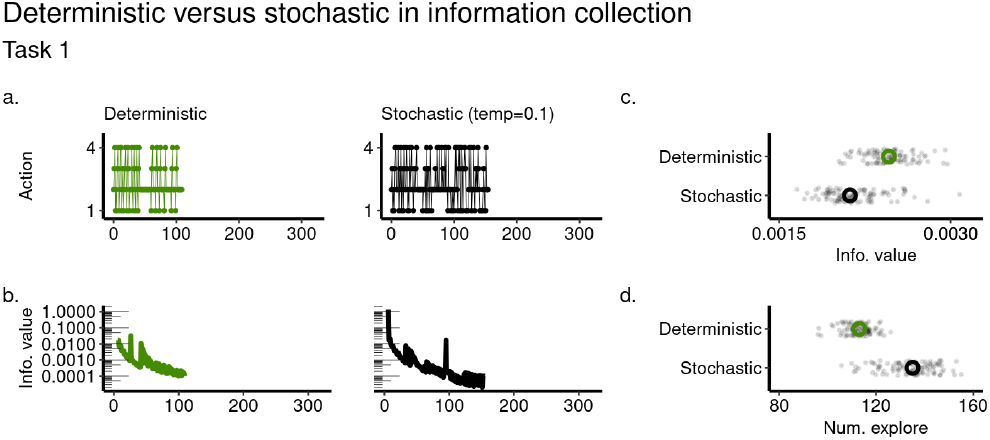
Comparing deterministic versus stochastic variations of the same curiosity algorithm in Task 1. Deterministic results are shown in the left column and stochastic results are shown in the right. **a**. Examples of exploration behavior. **b**. Information value plotted with time. Matches the behavior shown in a. Large values are prefered. **c**. Average information value for 100 independent simulations. er of steps it took to reach the stopping criterion in c., e.g. the boredom threshold *eta* (described below). Smaller numbers of steps imply a faster search.

### Reward collection - simulations

#### Bandits

Can curious search solve reward collection problems better than other approaches? To find out we measured the total reward collected in several bandit tasks (Figure 3b-g). We considered eight agents, including ours (Supplemental table S1). Each agent differed only in their exploration strategy. Examples of these strategies are shown in Figure. 5a. Note our union is denoted in green throughout. Without loss of generality, *E* is defined using a Bayesian information gain (14) which we sometimes abbreviate as “IG” (see Methods).

**Fig. 5.**
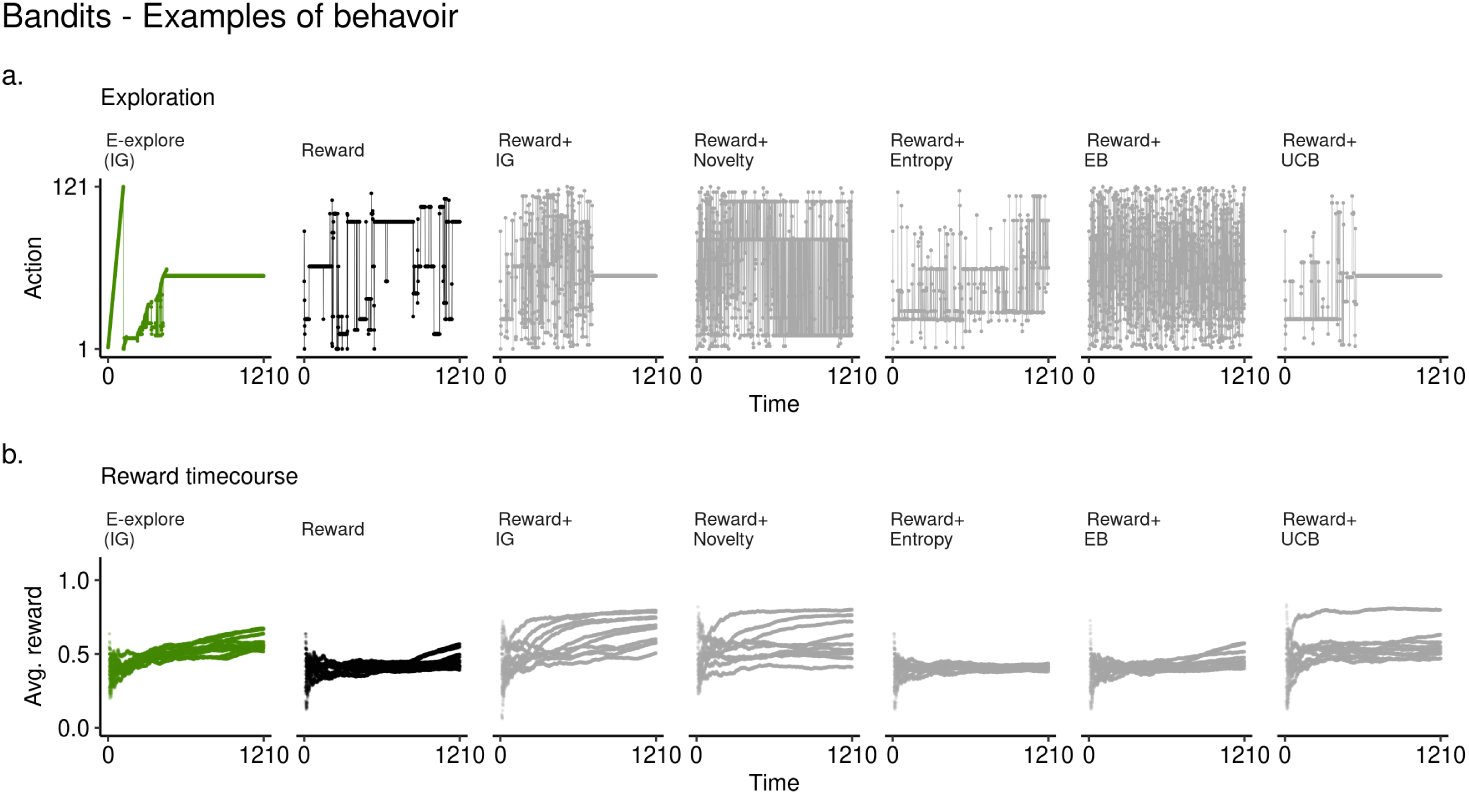
Behavior on a complex high-dimensional action space (Task 5). **a**. Examples of strategies for all agents (1 example). **b**. Average reward time courses for all agents (10 examples).

Despite E-exploration never optimizing for reward value, our method of pure exploration (69) matched or in some cases outperformed standard approaches that rely on reward value, at least in part.

In Figure 6 present overall performance in some standard bandit tasks: simple (Figure 6b), sparse (Figure 6c), high dimensional (Figure 6e) and non-stationary (Figure 6f). Overall, our approach matches or sometimes exceeds all the other exploration strategies (Figure 6a)).

**Fig. 6.**
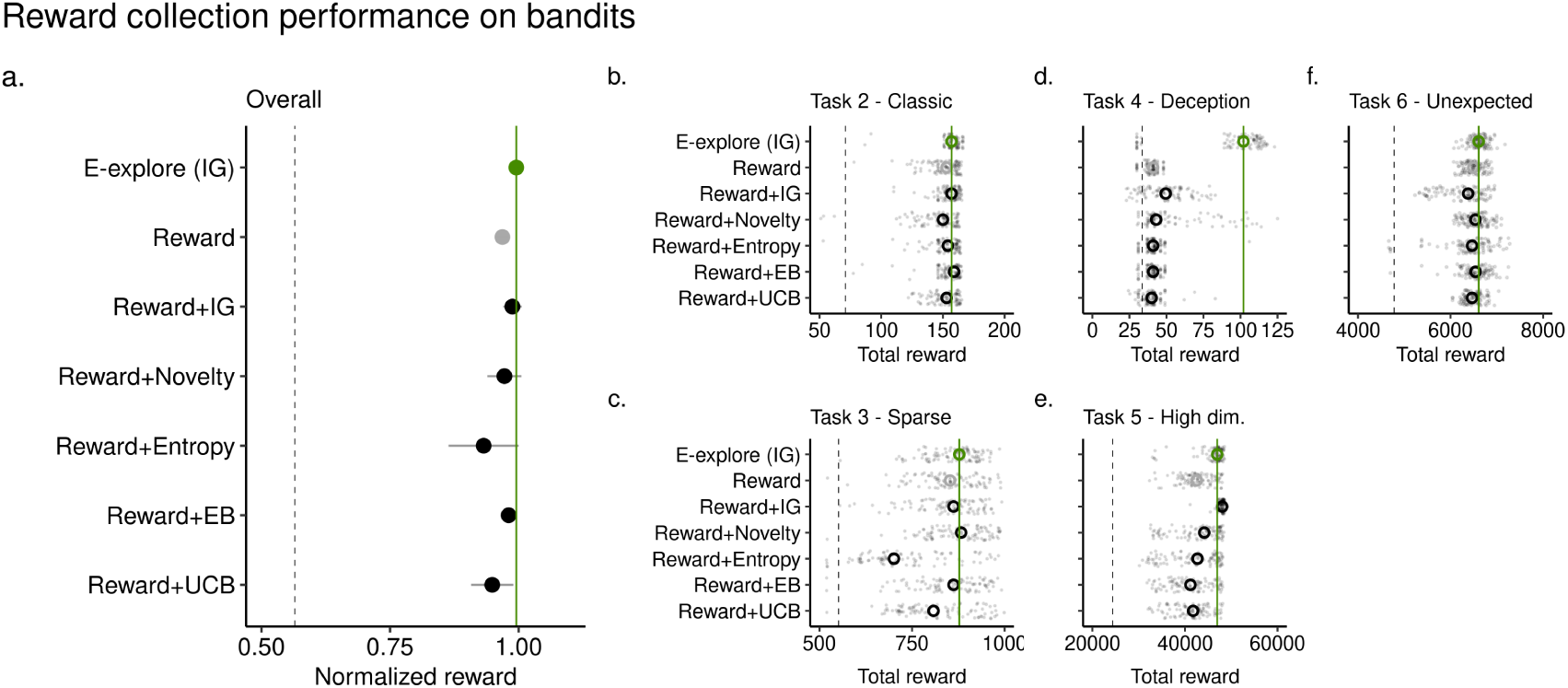
Reward collection performance. **b**. Overall results. Total reward collected for each of the four tasks was normalized. Dot represents the median value, error bars represent the median absolute deviation between tasks (MAD). **b**. Results for Task 2, which has four choices and one clear best choice. **c**. Results for Task 3, which has 10 choices and very sparse positive returns. **d**. Results for Task 4, whose best choice is initially “deceptive” in that it returns suboptimal reward value over the first 20 trials. **e**. Results for Task 6, which has 121 choices and a quite heterogeneous set of payouts but still with one best choice. **f**. Results for Task 7, which is identical to Task 6 except the best choice was changed the worst. Agents were pre-trained on Task 6, then tested on 7. In panels b-e, grey dots represent total reward collected for individual experiments while the large circles represent the experimental median.

With the exception of Task 4, performance improvements, when present, were small. This result might not look noteworthy. We argue it is because our exploration strategy never optimizes for reward value explicitly. Yet, we meet or exceed exploration strategies which do. In other words, when intrinsic curiosity is used in an optimal trade-off with extrinsic reward, curiosity does seem sufficient in practice.

In Figure 6f we considered deception. In this bandit there was an initial (misleading) 20 step decline in reward value (Figure 3d). On this task, other exploration strategies produced little better than chance performance. When deception is present, externally motivated exploration is a liability.

In Figure 7 we tested the robustness of all the agents to environmental mistunings. We re-examined total rewards collected across all model tunings or hyperparameters (Methods). Here E-explore was markedly better, with both substantially higher total reward when we integrate over all parameters (Figure 7a) and when we consider the tasks individually (Figure 7b-f).

**Fig. 7.**
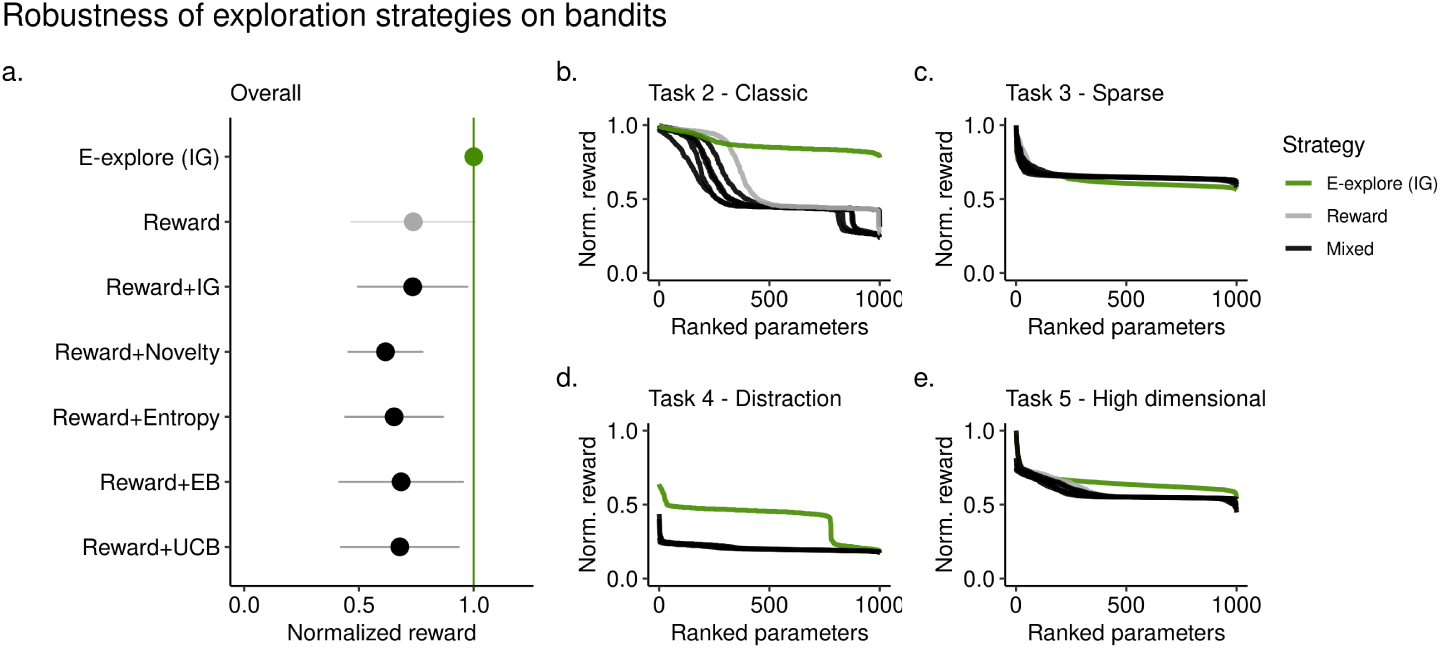
Exploration parameter sensitivity. **a**. Integrated total reward (normalized) across 1000 randomly chosen exploration hyperparameters. Dots represent the median. Error bars are the median absolute deviation. **b-f**. Normalized total reward for exploration 1000 randomly chosen parameters, ranked according to performance. Each subpanel shows performance on a different task. Lines are colored according to overall exploration strategy - E-explore (ours), reward only, or a mixed value approach blending reward and an exploration bonus).

#### Foraging

Bandit tasks are a simple means to study the explore-exploit problem space but in an abstract form. They do not adequately describe more natural settings where there is a physical space.

In Figure 8 we consider a foraging task defined on a 2d grid. In our foraging task for any position (*x, y*) there is a noisy “scent” which is “emitted” by each of 20 randomly placed reward targets. A map of example targets is shown in Figure 8a. The scent is a 2d Gaussian function centered at each target, corrupted by 1 standard deviation of noise (not shown).

**Fig. 8.**
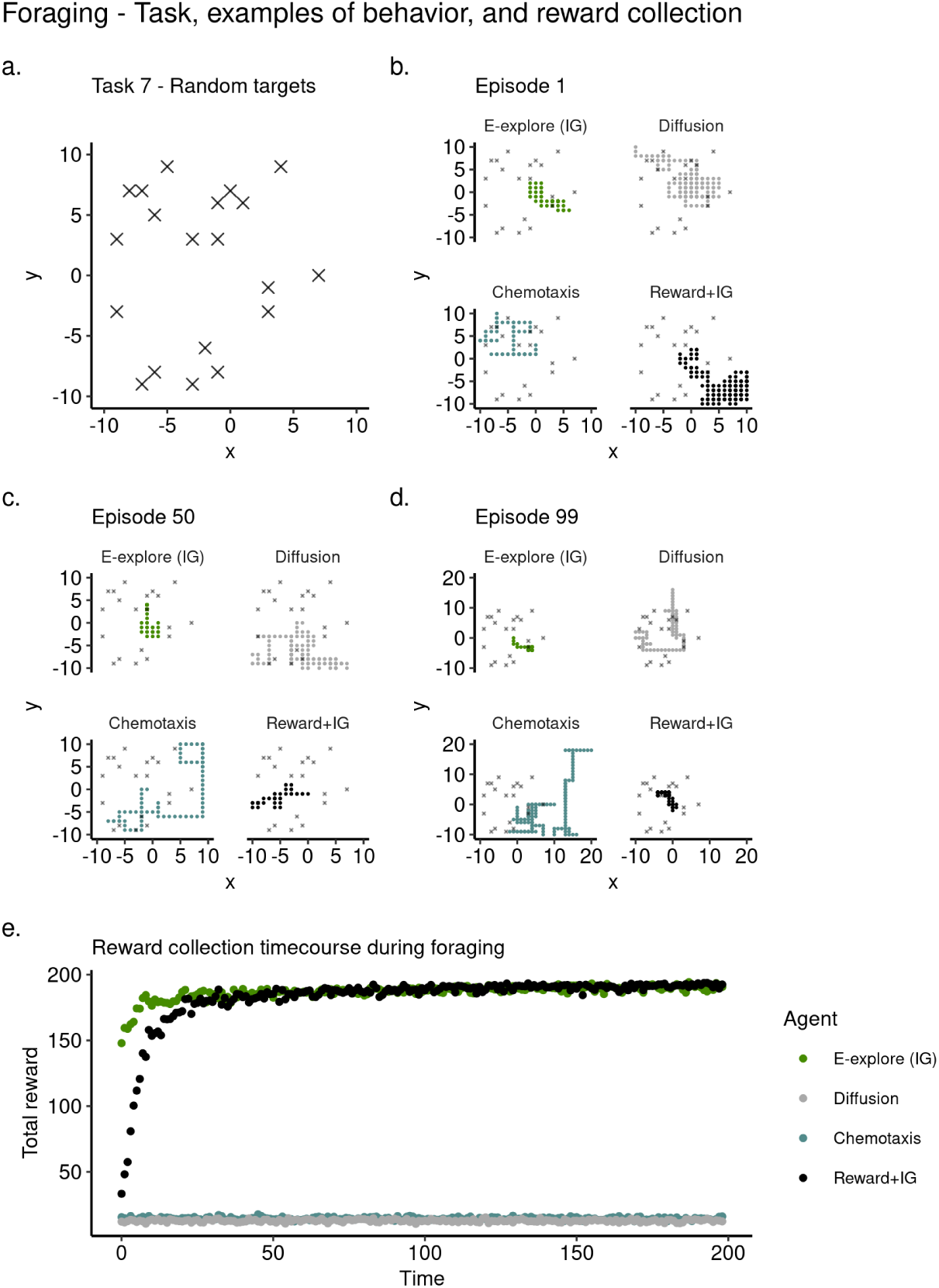
Reward collection during foraging (Task 8). See *Methods* for full details. **a**. Illustration of a 2 dimensional reward foraging task. **b-d**. Examples of agent behavior, observed over 200 time steps. a. Without any training. b. After 50 training episodes. c. After 99 training episodes. **e**. Total rewards collected over time, median value of 100 independent simulations.

In Figure 8b-d we show examples of foraging behavior by 4 agents (Supplemental table S2). Two of them are learning agents, ours and a mixed value reinforcement learning model. One is a chemotaxis agent. The other is a random search agent. Figure 8b shows foraging behavior before any learning. Figure 8c shows behavior halfway through the 100 experiments we considered. Figure 8d represents an example of the final behaviors. Notice how over this series ours, and the mixed value agent, learn to focus their foraging to a small targeted area as is best for this task.

In Figure 8e we show the reward acquisition time course averaged across all learning episodes. This measures both a learning progress and final task performance. Both reward learning agents plateau to the max value of 200, but ours (green) does so more quickly. In contrast, neither the chemotaxis or random agents show a change in profile. The takeaway for Task 8 is our method does scale well to 2d environments and can, at least in this one task, outperform standard approaches here as well.

## Discussion

We have shown that competitive union of pure curiosity with pure reward collection leads to a simple solution to the explore-exploit trade-off. Curiosity matches or exceeds reward-based strategies, in practice. That is, when rewards are sparse, high-dimensional, or non-stationary. It uniquely overcomes deception. Curiosity is also far more robust than the standard algorithms. We have derived a measure of information value via learning progress, but done in an axiomatic way it is possible to directly measure. This measure also properly generalizes many prior efforts to formalize curiosity.

In summary, we have used theory and simulations to argue scientists can safely set aside intuitions which suggest curious search is too inefficient to be generally practical. We have not shown our account can describe animal behavior in detail. For this, we have three strong experimental predictions to make.

1. **Invariant exploration strategy**. The presence of reward does not itself change the exploration strategy. For example, in (70) the exploration patterns of rats in a maze did change when a food reward was present at the maze’s end. *Auxiliary assumptions*. Learning about the environment is possible and there are some observations available to learn from.
2. **Deterministic search**. It is possible to predict exactly exploratory choices made during a curious search because the optimal search policy is strictly deterministic. To evaluate this we must move from population-level distribution analysis of behavior–which could be generated by bot deterministic or stochastic–to also examining moment-by-moment prediction of animal behavior (71, 72). *Auxiliary assumptions*: The noise in the neural circuits which implements behavior is sufficiently weak so it does not dominate the behavior itself.
3. **Well defined periods**. There should be well-defined periods of exploration and exploitation. For example, fitting a hidden markov model to the decision space should yield only two hidden states–one for exploration and one for exploitation. As in, (73).

### Questions and answers

We have been arguing that an established problem in decision theory–the classic dilemma–has been difficult or impossible to solve not because it must be, but because the field historically took a wrong view of exploration. We have also defined information value without considering the direct usefulness of the information learned. These are controversial viewpoints. So let’s consider some questions and answers.

#### Why curiosity?

Our use of curiosity rests on three well established facts. 1. Curiosity is a primary drive in most, if not all, animal behavior (15). 2. Curiosity is as strong, if not sometimes stronger, than the drive for reward (25, 74, 75). 3. Curiosity as an algorithm is highly effective at solving difficult optimization problems (9, 16, 27, 31, 32, 49, 58, 76, 77).

#### Is this a slight of hand, theoretically?

Yes. We have taken one problem that cannot be solved and replaced it with another related problem that can be. In this replacement we swap one behavior, extrinsic reward seeking, for another, curiosity.

#### Is this too complex?

Perhaps turning a single objective into two, as we have, is too complex an answer. If this is true, then it would mean our strategy is not a parsimonious solution to the dilemma. Should we reject it on that alone?

Questions about parsimony can be sometimes resolved by considering the benefits versus the costs of added complexity. The benefits are an optimal value solution to the exploration versus exploitation trade-off. A solution which seems especially robust to model-enviroment mismatch. At the same time curiosity-as-exploration can build a model of the environment (78), useful for later planning (79, 80), creativity, imagination (81), while also building diverse action strategies (32, 77, 82, 83).

#### Is this too simple?

So have we “cheated” by changing the dilemma problem in the way we have?

The truth is we might have cheated, in this sense: the dilemma might have to be as hard as it has seemed in the past. But the general case for curiosity is clear and backed up by the brute fact of its widespread presence in the animal kingdom. The question is: is curiosity so useful and so robust that it is sufficient for all exploration with learning (27).

The answer to this question is empirical. If our account does well in describing and predicting animal behavior, that would be some evidence for it (2, 29, 36, 74, 75, 84–88). If it predicts neural structures (89, 90), that would be some evidence for it. If this theory proves useful in machine learning, that would be some evidence for it (9, 27, 28, 31, 32, 46, 48, 82, 88, 91). In other words how simple, complex, or parsimonious an theoretical idea comes down to usefulness.

#### What about truth?

IIn other prototypical examples information value comes from prediction errors or is otherwise measured by how well learning corresponds to the environment (92–94) or how useful the information might be in the future (30). Colloquially, one might call this “truth seeking”.

As a people we pursue information which is fictional (95), or based on analogy (96), or outright wrong (97). Conspiracy theories, disinformation, and more mundane errors, are far too commonplace for information value to rest on mutual information or error. This does not mean that holding false beliefs cannot harm survival but this is a second order question as far as information value goes. The consequences of error on information value come after the fact.

#### What about information theory?

In short, the problems of communication of information and its value are wholy different issues that require different theories.

Weaver (98) in his classic introduction to the topic, describes information as a communication problem with three levels. **A**. The technical problem of transmitting accurately. **B**. The semantic problem of meaning. **C**. The effectiveness problem of using information to change behavior. He then describes how Shannon’s work addresses only the problem **A**., which let’s information theory find broad application.

Valuing information is not, at its core, a communications problem. It is a problem in personal history. Consider Bob and Alice who are having a phone conversation, and then Tom and Bob who have the exact same phone conversation. The personal history of Bob and Alice will determine what Bob learns in their phone call, and so determines what he values in that phone call. Ro so we argue. This might be different from what Bob learns when talking to Tom, even when the conversations was identical. What we mean to show by this example is that the personal history, also known as past memory, defines what is valuable on a channel.

We have summarized this diagrammatically, in Figure 9.

**Fig. 9.**
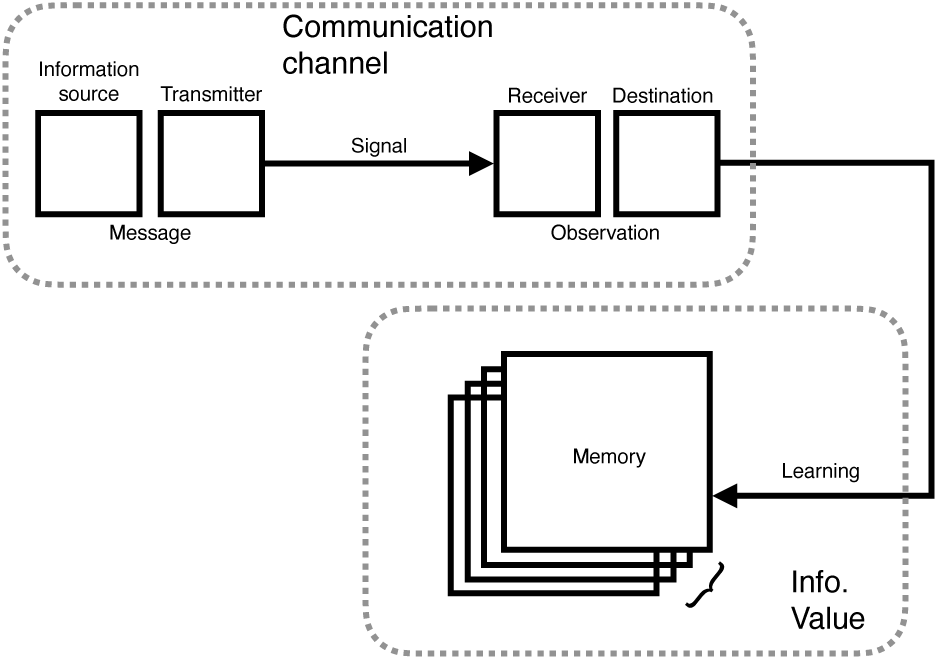
The relationship between the technical problem of communication with the technical problem of value. Note how value is not derived from the channel directly. Value comes from learning about observations taken from the channel, which in turn depends on memory.

Finally, we argue there is an analogous set of levels for information value as those Weaver describes for information. There is the technical problem of judging how much was learned. This is the one we address, like Shannon. There is the semantic problem of what this learning “means” and also what its consequences are. There is also the effectiveness problem of using what was learned to some other effect.

#### Does value as a technical problem even make sense?

Having a technical definition of information value, free from any meaning, might seem counterintuitive for any value measure. We suggest it is not any more or less counterintuitive than stripping information of meaning.

#### So you suppose there is always a positive value for learning of all fictions, disinformation, and deceptions?

We must. This is a most unexpected prediction. Yet sit seems consistent with the behavior of humans, and animals alike. Humans do consistently seek to learn falsehoods. Our axioms suggest why this is rational.

#### Was it necessary to build a general theory for information value to describe curiosity?

No. We made an idealistic choice that worked out. The field of curiosity studies has shown that there are many kinds of curiosity (9, 20, 25, 48, 49, 58, 74–76, 82, 85, 99–102). At the extreme limit of this diversity is a notion of curiosity defined for any kind of observation, and any kind of learning. This is what we offer. At this limit we can be sure to span fields, addressing the problem of information value as it appears in computer science, machine learning, game theory, psychology, neuroscience, biology, economics, among others.

#### What about other models of curiosity?

Curiosity has found many specific definitions (45, 75). Curiosity has been described as a prediction error, by learning progress (9, 103). Schmidhuber (9) noted the advantage of looking at the derivative of errors, rather than errors directly. Itti (104) and others (14, 20, 23, 105) have taken a statistical and Bayesian view often using the KL divergence to estimate information gain or Bayesian surprise. Other approaches have been based on adversarial learning, model disagreement (58), or model compression (20). Some measures focused on past experience (18, 78, 106). Others focused on future planning and imagination (107–109). Graphical models of memory have been useful (110).

What distinguishes our approach to information value is we focus on memory dynamics, and base value on some general axioms. We try to embody the idea of curiosity “as learning for learning’s sake”, not done for the sake of some particular goal (111), or for future utility (30). Our axioms were however designed with all these other definitions in mind. In other words, we do not aim to add a new metric. We aim to generalize the others.

#### Is information a reward?

If reward is any quantity that motivates behavior, then our definition of information value is a reward, an intrinsic rewaard. This last point does not mean that information value and environmental rewards are interchangeable however. Rewards from the environment are a conserved resource, information is not. For example, if a rat shares a potato chip with a cage-mate, it must break the chip up leaving it less food for itself. While if a student shares an idea with a classmate, that idea is not divided up. It depends, in other words.

#### But isn’t curiosity impractical?

It does seem curiosity can lead away from a needed solution as towards it. Consider children, who are the prototypical curious explorers (29, 74). This is why we focus on the derivatives of memory and limit curiosity with boredom, as well as counter curiosity with a drive for reward collecting (i.e., exploitation). All these elements combined seek to limit curiosity without compromising it.

Let’s consider colloquially how science and engineering can interact. Science is sometimes seen as an open-ended inquiry, whose goal is truth but whose practice is driven by learning progress, and engineering often seen as a specific target driven enterprise. They each have their own pursuits, in other words, but they also learn from each other often in alternating iterations. Their different objectives is what makes them such good long-term collaborators.

In a related view Gupta et al (1) encouraged managers in business organizations to strive for a balance in the exploitation of existing markets and ideas with exploration. They suggested managers should pursue periods of “punctuated equilibrium”, where employees work towards either pure market exploitation or pure curiosity-driven exploration to drive future innovation.

#### What is boredom, besides being a tunable parameter?

A more complete version of the theory would let us derive a useful or even optimal value for boredom if given a learning problem or environment. We cannot do this. It is the next problem we will work on, and it is important.

#### Does this mean you are hypothesizing that boredom is actively tuned?

Yes we are predicting that.

#### But can animals tune boredom?

Geana and Daw (61) showed this in a series of experiments. They reported that altering the expectations of future learning in turn alters self-reports of boredom. Others have shown how boredom and self-control interact to drive exploration (59, 60, 112).

#### Do you have any evidence in animal behavior and neural circuits?

There is some evidence for our theory of curiosity in psychology and neuroscience, however, in these fields curiosity and reinforcement learning have developed as separate disciplines (3, 74, 85). Indeed, we have highlighted how they are separate problems, with links to different basic needs: gathering resources to maintain physiological homeostasis (113, 114) and gathering information to decide what to learn and to plan for the future (3, 56). Here we suggest that though they are separate problems, they are problems that can, in large part, solve one another. This insight is the central idea to our view of the explore-exploit decisions.

Yet there are hints of this independent cooperation of curiosity and reinforcement learning out there. Cisek (2019) has traced the evolution of perception, cognition, and action circuits from the Metazoan to the modern age (89). The circuits for reward exploitation and observation-driven exploration appear to have evolved separately, and act competitively, exactly the model we suggest. In particular he notes that exploration circuits in early animals were closely tied to the primary sense organs (i.e. information) and had no input from the homeostatic circuits (89, 113, 114). This neural separation for independent circuits has been observed in some animals, including zebrafish (115) and monkeys (36, 116).

#### Is the algorithmic run time practical?

Computer scientists often study the run time of an algorithm as a measure of its efficiency. The worst case algorithmic run time of our method is linear and additive in the independent policies. If it takes *T*_*E*_ steps for *π*_*E*_ to converge, and *T*_*R*_ steps for *π*_*R*_, then the worst case run time for *π*_*ER*_ is *T*_*E*_ + *T*_*R*_.

## Methods and Materials

### Tasks

We studied seven bandit tasks. On each trial there were a set of *n* choices, and the agent should try and learn the best one. Each choice action returns a “payout” according to a predetermined probability. Payouts are information, reward, or both (Figure 3).

*Task 1* was designed to examine information foraging. There were no rewards. There were four choices. Three of these generated either a “yellow” or “blue” symbol, with a set probability. See Figure 3**a**.

*Task 2* was a simple simple bandit, designed to examine reward collection. At no point does the task generate information. Rewards were 0 or 1. There were four choices. The best choice had a payout of *p*(*R* = 1) = 0.8. This is a much higher average payout than the others (*p*(*R* = 1) = 0.2). See Figure 3**b**.

*Task 3* was designed with very sparse rewards (117, 118). There were 10 choices. The best choice had a payout of *p*(*R* = 1) = 0.02. The other nine had, *p*(*R* = 1) = 0.01 for all others. See Figure 3c.

*Task 4* had deceptive rewards. By deceptive we mean that the best long-term option presents itself initially with a lower value. The best choice had a payout of *p*(*R >* 0) = 0.6. The others had *p*(*R >* 0) = 0.4. Value for the best arm dips, then recovers. This is the “deception” It happens over the first 20 trials. Rewards were real numbers, between 0-1. See Figure 3d.

*Tasks 5-6* were designed with 121 choices, and a complex payout structure. Tasks of this size are at the limit of human performance (119). We first trained all agents on *Task 6*, whose payout can be seen in Figure3f-g. This task, like the others, had a single best payout *p*(*R* = 1) = 0.8. After training for this was complete, final scores were recorded as reported, and the agents were then challenged by *Task 7. Task 7* was identical except that the best option was changed to be the worst *p*(*R* = 0) = 0.2 (Figure 3f-g).

*Task 7* was a spatial foraging task (Figure 8a) where 20 renewing targets were randomly placed in a (20, 20) unit grid. Each target “emitted” a 2d Gaussian “scent” signal with a 1 standard deviation width. Agents began at the (0,0) center position and could move *(up, down, left, right)*. If the agent reached a target it would receive reward (1), continuously. All other positions generate 0 rewards. Odors from different targets were added.

### Agents

We considered two kinds of specialized agents. Those suited to solve bandit tasks and those suited to solve foraging tasks.

#### Bandit: E-explore

Our algorithm. It pursued either pure exploration based on *E* maximization or pure exploitation based on pure *R* maximization. All its actions are deterministic and greedy. Both *E* and *R* maximization was implemented as an actor-critic architecture (3) where value updates were made in crtic according to Eq. 19. Actor action selection was governed by,

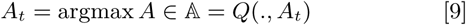

where we use the “.” to denote the fact our bandits have no meaningful state.

This agent has two parameters, the learning rate *α* (Eq. 19) and the boredom threshold *η* (Eq. 8). It’s payout function was simply, *G* = *R*_*t*_ (Eq. 19).

#### Bandit: Reward

An algorithm whose objective was to estimate the reward value of each action, and stochastically select the most valuable action in an effort to maximize total reward collected. It’s payout function was simply, *G* = *R*_*t*_ (Eq. 19). Similar to the E-explore agent, this agent used actor-critic, but its actions were sampled from a softmax / Boltzman distribution,

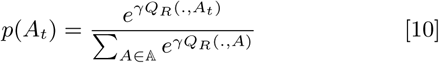

Where *γ* is the “temperature” parameter. Large *γ* generate more random actions. This agent has two parameters, the learning rate *α* (Eq. 19) and the temperature *γ >* 0 (Eq. 10). *Bandit: Reward+Info*. This algorithm was based on the Reward algorithm defined previously but it’s reward value was augmented or mixed with *E*. Specifically, the information gain formulation of *E* described below. It’s payout is, *G*_*t*_ = *R*_*t*_ + *βE*_*t*_ (Eq. 19). Critic values were updated using this *G*, and actions were selected according to Eq. 10. This agent has three parameters, the learning rate *α* (Eq. 19) the temperature *γ* (Eq. 10) and the exploitation weight *β >* 0. Larger values of *β* will tend to favor exploration.

#### Bandit: Reward+Novelty

This algorithm was based on the Reward algorithm defined previously but it’s reward value wass augmented or mixed with a novelty/exploration bonus (Eq. 11).

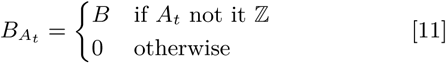

Where ℤ is the set of all actions that agent has taken so far in a simulation. Once all actions have been tried, *B* = 0.

This agent’s payout is, *G*_*t*_ = *R*_*t*_ + *B*_*t*_ (Eq. 19). Critic values were updated using this *G* and actions were selected according to Eq. 10. This agent has three parameters, the learning rate *α* (Eq. 19), the temperature *γ* (Eq. 10), and the bonus size *B >* 0.

#### Bandit: Reward+Entropy

This algorithm was based on the Reward algorithm defined previously but it’s reward value was augmented or mixed with an entropy bonus (Eq. 12). This bonus was inspired by the “softactor” method common to current agents in the related field of deep reinforcement learning (120).

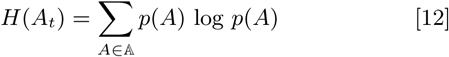

Where *p*(*A*) was estimated by a simple normalized action count.

This agent’s payout is, *G*_*t*_ = *R*_*t*_ + *βH*_*t*_ (Eq. 19). Critic values were updated using this *G*, and actions were selected according to Eq. 10. It has three free parameters, the learning rate *α* (Eq. 19), the temperature *γ* (Eq. 10), and the exploitation weight *β >* 0.

#### Bandit: Reward+EB

This algorithm was based on the Reward algorithm defined previously but it’s reward value was augmented or mixed with an evidence bound (EB) statistic (8) (Eq 13).

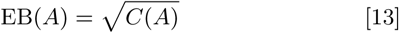

Where *C*(*A*) is a running count of each action, *A* ∈ 𝔸. This agent’s payout is, *G*_*t*_ = *R*_*t*_ + *β* EB (Eq. 19). Critic values were updated using this *G*, and actions were selected according to Eq. 14.

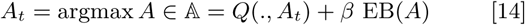

It has three two parameters, the learning rate *α* (Eq. 19) and the exploitation weight *β >* 0.

#### Bandit: Reward+UCB

This algorithm was based on the Reward algorithm defined previously, but it’s reward value was augmented or mixed with an upper confidence bound (UCB) statistic (8) (Eq 15).

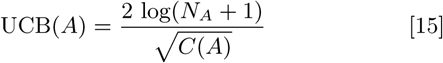

Where *C*(*A*) is a running count of each action, *A* ∈ 𝔸 and *N*_*A*_ is the total count of all actions taken. This agent’s payout is, *G*_*t*_ = *R*_*t*_ + *β* UCB (Eq. 19). Critic values were updated using this *G*, and actions were selected according to Eq. 16.

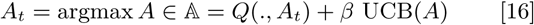

It has three two parameters, the learning rate *α* (Eq. 19) and the exploitation weight *β >* 0.

#### Forage: E-explore

This algorithm was identical to *Bandit: E-explore* except for the change in action space this task required (described above).

#### Forage: Reward+Info

This algorithm was identical to *Bandit: Reward+Info* except for change in action space the forage task required (as described in the *Task 8* section above).

#### Forage: Diffusion

This algorithm was designed to ape 2d brownian motion, and therefore to implement a form of random search (121). At every timestep this agent either continued in motion, if it had not traversed a previously sampled length *l*, or it drew a new *l* and random uniform direction, *d*_*f*_ = *U* (up, down, left, right). Movement lengths were sampled from an exponential distribution (Eq. 17).

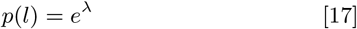

Where *λ* is the length scale which was set so *λ* = 1 in all experiments.

#### Forage: Chemotaxis

Movement in this algorithm was based on the Diffusion model, except it used a biased approach to decide when to turn. Specifically, it used the scent concentration gradient to bias its change in direction (Eq. 18).

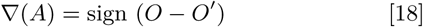

Where *O* is the sense or scent observation made by the agent at its current grid position, and sign returns 1 if the argument is positive, and −1 otherwise.

Turning decisions were based on a biased random walk model, often used in -taxis behaviors (122). In this there are two independent probability thresholds, *p*_*f*_ and *p*_*r*_. If the gradient is positive, then the *p*_*f*_ is “selected” and a coin flip happens, generating a random value 0 ≤ *x*_*i*_ ≤ 1. If *p*_*f*_ ≥ *x*_*i*_, then the current direction *d*_*f*_ is maintained, otherwise a new direction is randomly chosen (see Diffusion above for details). If the gradient was negative, then the criterion *p*_*r*_ ≥ *x*_*i*_ is used.

### Task memory for E-explore

In the majority of simulations we have used a Bayesian or Info Gain formulation for curiosity and *E*. Each task’s observations fit a simple discrete probabilistic model, with a memory “slot” for each action for each state. Specifically, probabilities were tabulated on state reward tuples, (*S, R*). To measure distances in this memory space we used the Kullback–Leibler divergence (20, 104, 123–126).

### Learning equations

Reward and information value learning for all agents on the bandit tasks were made using update rule below,

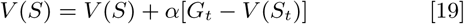

Where *V* (*S*) is the value for each state, *G*_*t*_ is the *return* for the current trial, either *R*_*t*_ or *E*_*t*_, and *α* is the learning rate (0 − 1]. See the *Hyperparameter optimization* section for information on how *α* is chosen for each agent and task.

Value learning updates for all relevant agents in the foraging task were made using the TD(0) learning rule (3),

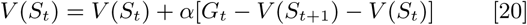

We assume an animal will have a uniform prior over the possible actions 𝔸 and set accordingly, *E*_0_ = Σ_*K*_ *p*(*A*_*k*_) log *p*(*A*_*k*_).

### Hyperparameter optimization

The hyperparameters for each agent were tuned independently for each task by random search (127). Generally, we reported results for the top 10 values, sampled from 1000 possibilities. Top 10 parameters and search ranges for all agents are shown in Supplemental Table S3. Parameter ranges were held fixed for each agent across all the bandit tasks.

## Acknowledgments

We wish to thank Jack Burgess, Matt Clapp, Kyle “Donovank” Dunovan, Richard Gao, Roberta Klatzky, Jayanth Koushik, Alp Muyesser, Jonathan Rubin, and Rachel Storer for their comments on earlier drafts. We also wishes to thank Richard Grant for his illustration work in Figure 1. The research was sponsored by the Air Force Research Laboratory (AFRL/AFOSR) award FA9550-18-1-0251. The views and conclusions contained in this document are those of the authors and should not be interpreted as representing the official policies, either expressed or implied, of the Army Research Laboratory or the U.S. government. TV was supported by the Pennsylvania Department of Health Formula Award SAP4100062201, and National Science Foundation CAREER Award 1351748.

## Mathematical Appendix

### Optimal substructure in memory

To find an optimal value solution for *E* using the Bellman equation we must prove our memory **M** has optimal substructure. This is because the normal route to a Bellman solution, which assumes the problem rests in a Markov Space, is closed to us. To understand why it is closed to us let’s consider a normal Markov case.

In Markov spaces there are a set of states where the transition to the next *S*_*t*_ depends only on the previous state *S*_*t*−1_. If we were to optimize over these states, as in typical reinforcement learning, we can know each transition as its own “subproblem”, dynamics programming approaches apply.

The problem for our definition of information value is that it relies on a memory definition which is necessarily composed of many past observations, arbitrarily, and so it cannot for certain be a Markov space. In fact, it is probably now. So if we wish to use a dynamics programming approach we need to find another way to establish optimal substructure, which is the ability to make the simpler independent “subproblems” dynamic programming and the Bellman equation rely on. Establishing these subproblems is the focus of Theorem 1, which proceeds by contradiction.

The “heavy lifting” in the proof is done by assuming we have a perfect forgetting function for the most recent past observation. This is what we denote asas *f* ^−1^. In this theorem we assume 𝒳, *A*, **M**, *f*, and 𝒯 are implicitly given.

In the theorem below which shows the optimal substructure of **M** we assume 𝒳, and *f* are given,

#### Theorem 1 (Optimal substructure).

If 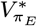 *is the optimal information value given by policy π*_*E*_, *a memory* **M**_*t*_ *has optimal substructure if the last observation S can be removed from* **M**, *by* **M**_*t*−1_ = *f* ^−1^*f* (𝒳, **M**_*t*_) *such that the resulting value* 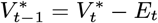 *is also optimal*.

*Proof*. Given a known optimal value *V* * given by *π*_*E*_ we assume for the sake of contradiction there also exists an alternative policy 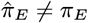 at time *t* that gives a memory 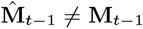 and for which 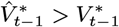.

To recover the known optimal memory **M**_*t*_ we lift 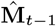 to 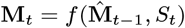. This implies 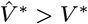 which contradicts the purported optimality of *V*\* and therefore *π*_*E*_. □

**Table S1.** Exploration strategies for bandit agents.

| Name | Class | Exploration strategy |
| --- | --- | --- |
| E-explore | Info. val. | Deterministic max. of information value |
| Random | Random | Random exploration |
| Reward | Mixed | Softmax sampling of reward |
| Reward+Info. | Mixed | Random sampling of reward + $\beta$ information value |
| Reward+Novelty | Mixed | Random sampling of reward + $\beta$ novelty signal |
| Reward+Entropy | Mixed | Random sampling of reward + $\beta$ action entropy |
| Reward+EB | Mixed | Random sampling of reward + $\beta$ visit counts |
| Reward+UCB | Mixed | Random sampling of reward + $\beta$ visit counts |

**Table S2.** Exploration strategies for foraging agents.

| Name | Class | Exploration strategy |
| --- | --- | --- |
| E-explore | Info. val. | Deterministic max. of information value |
| Diffusion | Random | Random exploration |
| Chemotaxis | Obs. | Sense gradient + Random exploration |
| Reward+Info. | Mixed | Random sampling of reward + $\beta$ information |

**Table S3.** Hyperparameter tuning - parameters and ranges.

| Task | Agent | Parameter | Range |
| --- | --- | --- | --- |
| Bandit | E-explore | $\eta$ | (1e-9, 1e-2) |
| Bandit | E-explore | $\alpha$ | (0.001, 0.5) |
| Forage | E-explore | $\eta$ | (1e-9, 1e-2) |
| Forage | E-explore | $\alpha$ | (0.001, 0.5) |
| Bandit | Reward | $\gamma$ | (0.001, 1000) |
| Bandit | Reward | $\alpha$ | (0.001, 0.5) |
| Bandit | Reward+Info. | $\gamma$ | (0.001, 1000) |
| Bandit | Reward+Info. | $\beta$ | (0.001, 10) |
| Bandit | Reward+Info. | $\alpha$ | (0.001, 0.5) |
| Forage | Reward+Info. | $\gamma$ | (0.001, 1000) |
| Forage | Reward+Info. | $\beta$ | (0.001, 10) |
| Forage | Reward+Info. | $\alpha$ | (0.001, 0.5) |
| Bandit | Reward+Novelty | $\gamma$ | (0.001, 1000) |
| Bandit | Reward+Novelty | $B$ | (1, 100) |
| Bandit | Reward+Novelty | $\alpha$ | (0.001, 0.5) |
| Bandit | Reward+Entropy | $\gamma$ | (0.001, 1000) |
| Bandit | Reward+Entropy | $\beta$ | (0.001, 10) |
| Bandit | Reward+Entropy | $\alpha$ | (0.001, 0.5) |
| Bandit | Reward+EB | $\beta$ | (0.001, 10) |
| Bandit | Reward+EB | $\alpha$ | (0.001, 0.5) |
| Bandit | Reward+UCB | $\beta$ | (0.001, 10) |
| Bandit | Reward+UCB | $\alpha$ | (0.001, 0.5) |

